# A Genome-based Model to Predict the Virulence of *Pseudomonas aeruginosa* Isolates

**DOI:** 10.1101/2020.06.09.143610

**Authors:** Nathan B. Pincus, Egon A. Ozer, Jonathan P. Allen, Marcus Nguyen, James J. Davis, Deborah R. Winter, Chih-Hsien Chuang, Cheng-Hsun Chiu, Laura Zamorano, Antonio Oliver, Alan R. Hauser

**Affiliations:** Department of Microbiology-Immunology, Northwestern University Feinberg School of Medicine, Chicago, Illinois, USA; Department of Medicine, Division of Infectious Diseases, Northwestern University Feinberg School of Medicine, Chicago, Illinois USA; University of Chicago Consortium for Advanced Science and Engineering, University of Chicago, Chicago, Illinois USA; Division of Data Science and Learning, Argonne National Laboratory, Argonne, Illinois, USA; Northwestern-Argonne Institute of Science and Engineering, Evanston, Illinois, USA; Department of Medicine, Division of Rheumatology, Northwestern University Feinberg School of Medicine, Chicago, USA; School of Medicine, College of Medicine, Fu-Jen Catholic University, New Taipei, Taiwan; Molecular Infectious Disease Research Center, Chang Gung Memorial Hospital, Chang Gung University, Taoyuan, Taiwan; Servicio de Microbiología y Unidad de Investigación, Hospital Universitari Son Espases, Institut d’Investigació Sanitaria Illes Balears, Palma de Mallorca, Spain

## Abstract

Variation in the genome of *Pseudomonas aeruginosa*, an important pathogen, can have dramatic impacts on the bacterium’s ability to cause disease. We therefore asked whether it was possible to predict the virulence of *P. aeruginosa* isolates based upon their genomic content. We applied a machine learning approach to a genetically and phenotypically diverse collection of 115 clinical *P. aeruginosa* isolates using genomic information and corresponding virulence phenotypes in a mouse model of bacteremia. We defined the accessory genome of these isolates through the presence or absence of accessory genomic elements (AGEs), sequences present in some strains but not others. Machine learning models trained using AGEs were predictive of virulence, with a mean nested cross-validation accuracy of 75% using the random forest algorithm. However, individual AGEs did not have a large influence on the algorithm’s performance, suggesting instead that the virulence prediction derives from a diffuse genomic signature. These results were validated with an independent test set of 25 *P. aeruginosa* isolates whose virulence was predicted with 72% accuracy. Machine learning models trained using core genome single nucleotide variants and whole genome k-mers also predicted virulence. Our findings are a proof of concept for the use of bacterial genomes to predict pathogenicity in *P. aeruginosa* and highlight the potential of this approach for predicting patient outcomes.

**IMPORTANCE:** *Pseudomonas aeruginosa* is a clinically important gram-negative opportunistic pathogen. As a species, *P. aeruginosa* has a large degree of heterogeneity both through variation in sequences found throughout the species (core genome) and the presence or absence of sequences in different isolates (accessory genome). *P. aeruginosa* isolates also differ markedly in their ability to cause disease. In this study, we used machine learning to predict the virulence level of *P. aeruginosa* isolates in a mouse bacteremia model based on genomic content. We show that both the accessory and core genome are predictive of virulence. This study provides a machine learning framework to investigate relationships between bacterial genomes and complex phenotypes such as virulence.

## INTRODUCTION

*Pseudomonas aeruginosa* is a ubiquitous gram-negative opportunistic pathogen that can infect a variety of hosts. Its ability to cause severe acute infections in susceptible patients and chronic infections in individuals with cystic fibrosis, coupled with increasing rates of antimicrobial resistance, make it an organism of particular concern to the medical community^1-3^. The *P. aeruginosa* species, however, is not monolithic but rather shows a large amount of genomic diversity both through polymorphisms and differences in gene content^4-6^. As routine whole genome sequencing becomes increasingly feasible, understanding how these genomic differences impact the pathogenicity of *P. aeruginosa* may allow clinicians to rapidly identify high-risk infections and researchers to select the most high-yield strains for further study.

As with other bacteria, the genome of *P. aeruginosa* can be divided into a core genome, made up of sequences common to the species, and an accessory genome, made up of sequences present in some strains but not others^6,7^. While only 10-15% of a typical strain’s genome is accessory, these sequences when combined from all strains comprise the vast majority of the *P. aeruginosa* pangenome^4,7,8^. Variations in both the core and accessory genomes play an important role in the virulence of any given *P. aeruginosa* strain. Core genome mutations that accumulate in *P. aeruginosa* strains during chronic infection of cystic fibrosis patients lead to decreased markers of *in vitro* virulence^9^, and these strains have attenuated virulence in animal models of acute infection^10^. Genomic islands, major components of the accessory genome, are enriched for predicted virulence factors^11^. Several genomic islands in *P. aeruginosa*, including those containing the type III secretion system (T3SS) effector gene *exoU*, have been shown to enhance pathogenicity in multiple infection models^12-14^. We recently mined the accessory genome and identified multiple novel factors important in virulence in a mouse model of bacteremia^15^. Conversely, a study using a *Caenorhabditis elegans* model identified several *P. aeruginosa* accessory genes whose presence reduced virulence^16^. Further, the presence of active CRISPR systems was associated with increased virulence^16^, supporting the hypothesis that many horizontally transferred elements are genetic parasites with respect to the host bacterium^17^. Because of its role in both increasing and decreasing the pathogenicity of individual *P. aeruginosa* strains, the accessory genome may serve as a useful predictor of an isolate’s virulence. This is not necessarily as simple as detecting individual virulence or anti-virulence factors. For example, *exoU* is a recognized virulence factor whose disruption dramatically attenuates a strain’s ability to cause disease^18,19^, but some strains naturally lacking *exoU* are more virulent than those possessing the gene^15^. As virulence is a complex and combinatorial phenotype, the strategy taken to study it must be appropriately robust to that complexity.

Machine learning is an increasingly important tool in bacterial genomics and has been extensively applied to the prediction of antimicrobial resistance and identification of potential resistance determinants. This has proven successful in a variety of species and using a variety of genomic features^20-26^. These studies have benefited from readily available whole genome sequencing and resistance data, as well as an often easily explainable phenotype. Researchers have also begun to apply machine learning techniques to predict bacterial pathogenicity. Examples include using discriminatory single nucleotide variants (SNVs) to predict *Staphylococcus aureus in vitro* cytotoxicity^27^, using variation in core genome loci to predict patient mortality in specific *S. aureus* clones^28^, and using predicted perturbations in protein coding sequences to classify *Salmonella* strains as causing either gastrointestinal or extraintestinal infections^29^. A support vector machine approach has been used to distinguish the transcriptomes of *P. aeruginosa* in human infection compared to *in vitro* growth^30^. However, to our knowledge there has been no study directly modeling *P. aeruginosa* pathogenicity from genomic content.

In this study, we utilized a machine learning approach to predict *P. aeruginosa* virulence in a mouse model of bloodstream infection based on genomic content. We found that there is signal within the accessory genome predictive of virulence, a finding validated using an independent test set of isolates. This appears to be through the detection of a diffuse genetic fingerprint rather than individual virulence or anti-virulence genes. The core genome also showed predictive signal for virulence.

## RESULTS

### Genomic and Virulence Characterization of *P. aeruginosa* Strains

To assess the ability of the *P. aeruginosa* genome to predict a given isolate’s virulence, we needed a large number of *P. aeruginosa* isolates with known whole genome sequences and *in vivo* virulence data. For this purpose, we used two previously reported collections: 98 archived isolates from adults with bacteremia at Northwestern Memorial Hospital (NMH) in Chicago, USA^31^ and 17 isolates from children with Shanghai fever, a *P. aeruginosa* infection presenting with sepsis and gastrointestinal symptoms, at Chang Gung Children’s Hospital in Taiwan^32^ (Supplementary Table 1). Together these formed a training set of 115 isolates. We performed whole genome sequencing for each of the isolates that had not been previously sequenced. Likewise, previously reported virulence data^15,32^ were supplemented with new experiments (Supplementary Table 2) to approximate the colony forming units (CFU) of each bacterial isolate necessary to cause pre-lethal illness in 50% of mice using a bacteremia model. From these data, a modified 50% lethal dose (mLD_50_) was estimated for each of the 115 *P. aeruginosa* isolates (Supplementary Table 3). The isolates showed a median mLD_50_ of 6.9 log_10_ CFU but with a wide range of pathogenicity in mice, varying by over 100-fold in the dose required to cause severe disease, as was previously reported for the NMH isolates^15^. For the purpose of this study, we classified those isolates with an estimated mLD_50_ below the median value for the group as “high virulence” and the remainder as “low virulence” (Figure 1A). These results provided a large collection of *P. aeruginosa* isolates with known whole genome sequences and virulence in a mouse bacteremia model.

**Figure 1.**
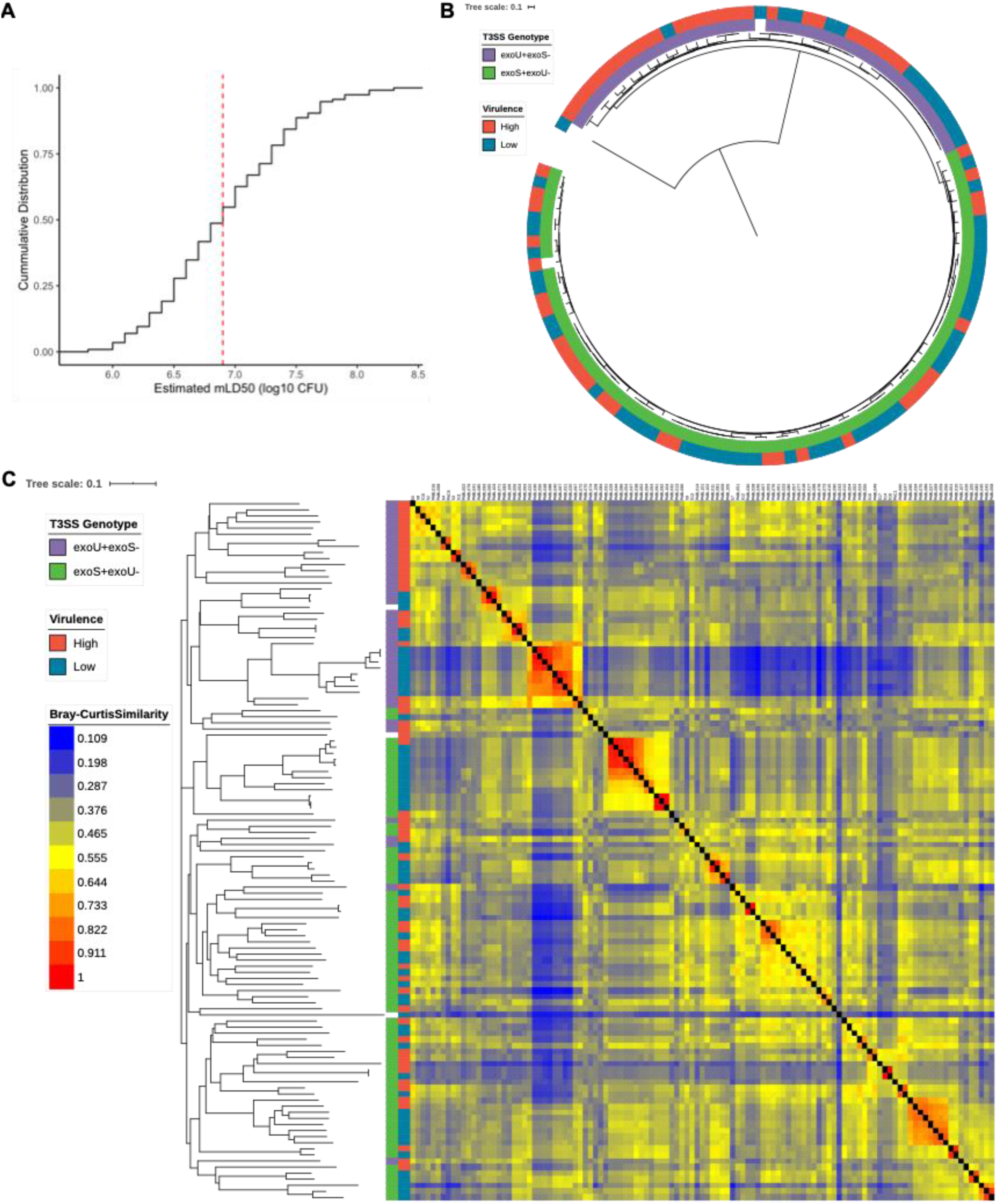
Virulence and genomic characteristics of the training set of 115 *P. aeruginosa* isolates. (A) Cumulative density function of estimated mLD_50_s for the 115 isolates in a mouse model of bacteremia. Isolates with estimated mLD_50_s less than the median value (red dashed line) were designated as high virulence, with the remainder designated as low virulence. (B) Mid-point rooted core genome phylogenetic tree of the 115 training isolates constructed from SNV loci present in at least 95% of genomes, annotated with T3SS genotype and virulence level. (C) Bray-Curtis dissimilarity heatmap comparing AGE presence in the 115 training isolates, weighted by AGE length, and accompanying neighbor joining tree. Isolates are annotated (from left to right) by T3SS genotype, virulence level, and the dissimilarity heatmap. A higher value indicates that two isolates have more similar accessory genomes.

We performed a phylogenenomic analysis to assess the diversity of the core genomes of all 115 isolates in the training set (Figure 1B). The core genome phylogenetic tree showed that the isolates are largely nonclonal and were found in both major clades of the species, which are mainly differentiable by the near-mutually exclusive presence of the T3SS effector genes *exoS* or *exoU*^*4,5*^. One distinct outlier isolate from the PA7-like clade was also present in the collection^4^. The *exoU+* clade contained a larger proportion of highly virulent isolates than the *exoS*+ clade. Although some clusters of closely related isolates shared the same virulence class, both major clades contained high- and low-virulence isolates.

We next defined the accessory genome of each of the 115 isolates in the training set. The accessory genome can be divided into accessory genomic elements (AGEs), discrete sequences found in the genomes of some isolates but not others^7^. For the purpose of this study, perfectly correlated sets of accessory sequences were grouped and considered as a single AGE. This means that noncontiguous sequences are treated as a single element if they are present and absent from the same isolates in the training set. Sets of accessory sequences totaling less than 200 bp were excluded from analysis. Using this approach, a total of 3,013 AGEs, with mean length 4,059 bp, median length 672 bp, and forming a pan-accessory genome of 12.2 Mb, were identified in these isolates (Supplementary Table 4). A Bray-Curtis dissimilarity heatmap of AGE presence/absence, weighted by the length of each AGE, shows that there is considerable accessory genomic variability in our collection (Fig. 1C). Consistent with previous findings^4^, the clade containing *exoS* and the clade containing *exoU* largely separate based on accessory genomic content, as evidenced by both Bray-Curtis dissimilarity and multiple correspondence analysis. Similar to the core genome phylogenetic analysis, some clusters of isolates with similar accessory genomes share a virulence rank, but both high and low virulence isolates show diverse AGE content (Figure 1C and Supplementary Figure 1A and B).

### Evaluating Machine Learning Models Predicting *P. aeruginosa* Virulence Based on Accessory Genome Content

We hypothesized that as the *P. aeruginosa* accessory genome is variable between strains^6,7,33^ and includes multiple known virulence determinants^12,13,15^, it would contain information predictive of strain virulence in mice. To test this hypothesis, we took a 10-fold nested cross-validation approach (Supplementary Figure 2). Using this approach, we tested the performance of four commonly used machine learning algorithms: random forest, l2-regularized logistic regression, elastic net logistic regression, and support vector classifier. Accessory genome content, in the form of AGE presence/absence, was used as features and virulence level (high or low) was used as labels during modeling. Nested cross-validation does not return a final machine learning model, instead examining how multiple models (with hyperparameters tuned through grid-search cross-validation) perform against held-out data. This process provides an estimate of how well a model trained through a given strategy will generalize to new data. All four algorithms performed similarly, with mean nested cross-validation accuracies of 0.75 (95% CI 0.69-0.80) for random forest, 0.75 (95% CI 0.65-0.85) for l2-regularized logistic regression, 0.72 (95% CI 0.65-0.79) for elastic net logistic regression, and 0.74 (95% CI 0.67-0.81) for support vector classifier. Other performance metrics showed similar ranges of values (Figure 2). Notably, the accuracy of all four algorithms was substantially higher than the null accuracy of simply predicting all isolates to be the majority class, which in this case was the prevalence of low virulence isolates (0.51). This indicates that there is signal in the accessory genome predictive of virulence in *P. aeruginosa*. Since all four machine learning algorithms performed similarly in nested cross-validation, we chose the random forest approach for further investigation.

**Figure 2.**
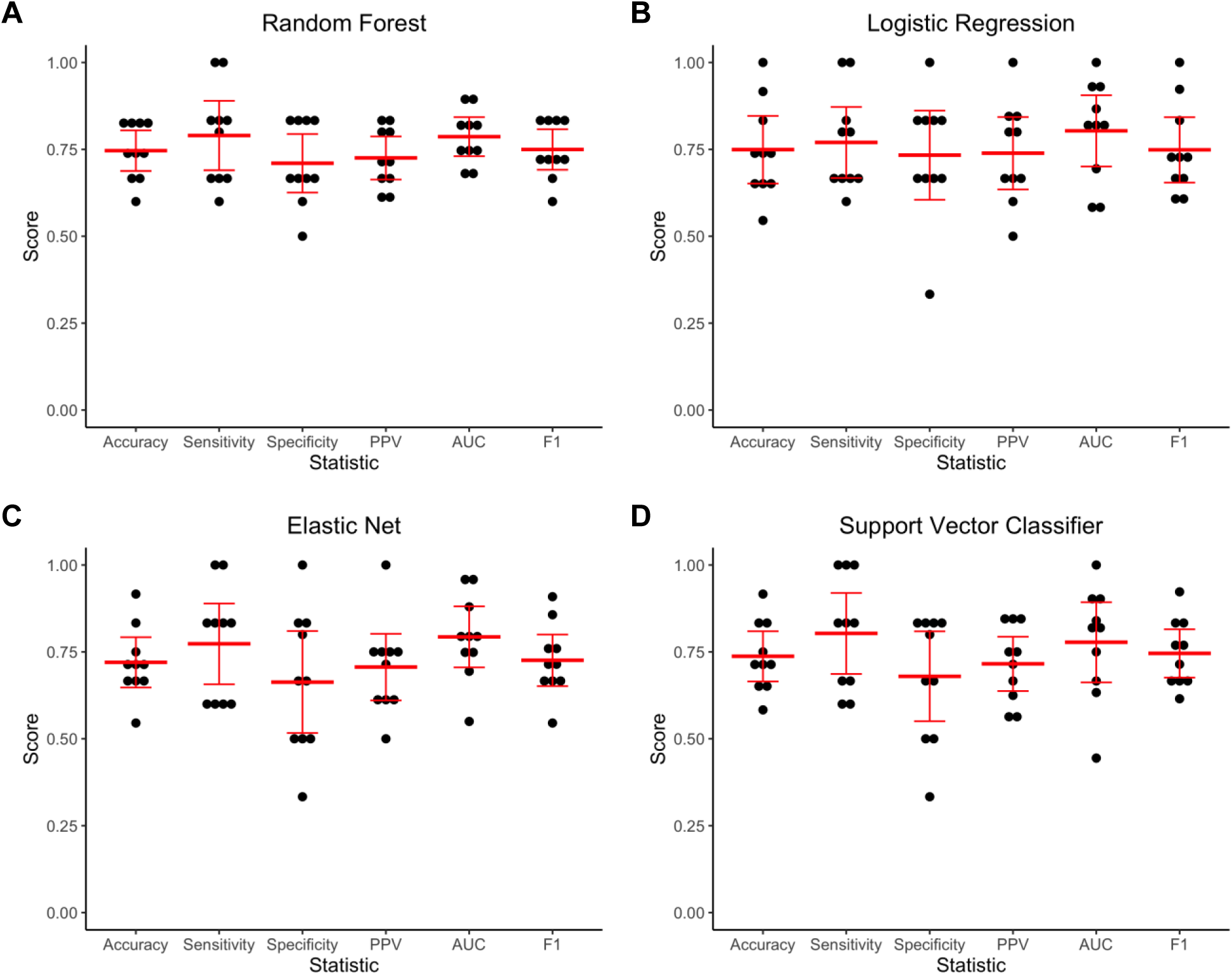
Nested 10-fold cross-validation performance of machine learning algorithms in predicting *P. aeruginosa* virulence in mice based on accessory genomic content. (A) Random forest, (B) l2-regularized logistic regression, (C) elastic net logistic regression, and (D) support vector classifier algorithms were tested. Accuracy, sensitivity, specificity, positive predictive value (PPV), area under the receiver operating characteristic curve (AUC), and F1 score were determined for each cross-validation fold (black dots). The mean and 95% confidence interval of each statistic are indicated in red.

We next evaluated whether sample size limited the performance of the random forest approach. We tested how accuracy of a model changed with increasing training set size, both against training and cross-validation examples (Figure 3A). While the training and cross-validation performance for the random forest model did not completely converge as more training examples were added, the learning curve showed that we are unlikely to see substantial improvement in cross-validation accuracy with additional training isolates. A caveat to this result is that the learning curve assumes our training set is adequately representative of *P. aeruginosa* genetic diversity.

**Figure 3.**
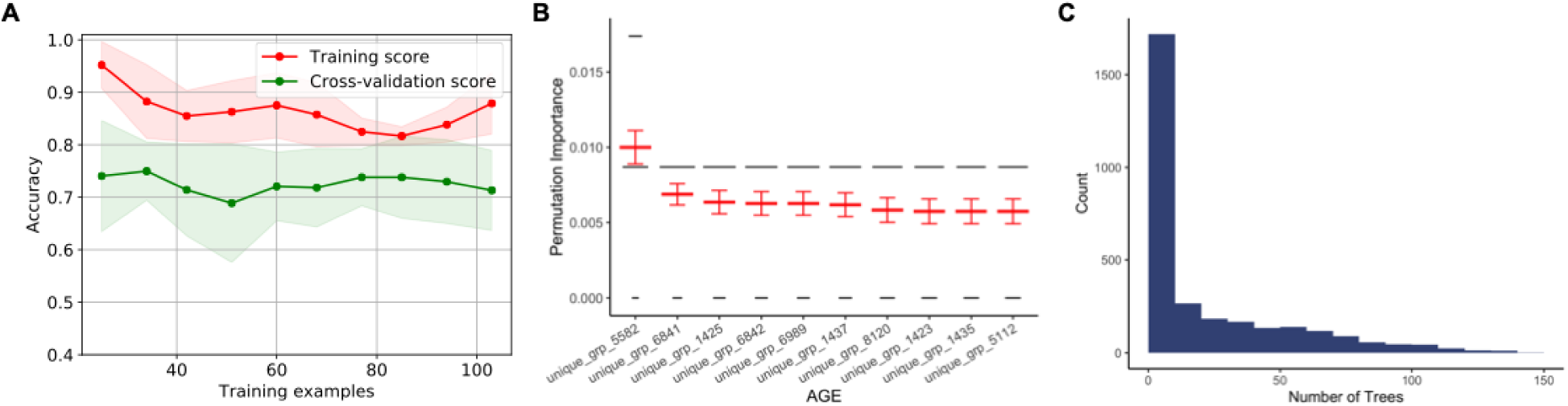
Evaluation of the random forest algorithm in predicting *P. aeruginosa* virulence based on accessory genomic content. (A) Learning curve showing change in mean training accuracy (red line) and cross validation accuracy (green line) in predicting *P. aeruginosa* virulence as increasing numbers of isolates are used to train the random forest model. Shading indicates the 95% confidence interval. Assessments at each number of training examples were through 10-fold nested cross-validation. (B) Out-of-bag permutation importance for the 10 most important AGEs in the random forest model, showing decrease in accuracy when these AGEs were randomly permuted. Permutation importance testing was performed 100 times, with the results of each test represented by the width of the black lines and the mean and 95% confidence interval indicated in red for each AGE. (C) Histogram indicating how many trees within the random forest model contained each AGE (feature), out of a total of 10,000 trees.

To further probe the characteristics of the random forest approach, we built a final random forest model using all 115 isolates in the training set. The out-of-bag accuracy (performance on the out-of-bag samples not included in each of the 10,000 decision trees making up the random forest) of this model was 0.75 (Supplementary Table 6), which is consistent with our nested cross-validation results. When assessed against the training isolates, the model showed an accuracy of 0.79, consistent with the trend in training accuracies observed in the learning curve (Supplementary Table 6 and Figure 3A). The training accuracy can be thought of as an idealized maximal performance and supports the conclusion that additional training examples are unlikely to substantially improve the model.

We next investigated which AGEs were most critical in making a prediction of high or low virulence in this model. We calculated the permutation importance (the mean decrease in model accuracy if a given feature is randomly permuted) for each AGE. To do this, we randomly permuted each AGE 100 times and the determined the impact on out-of-bag accuracy. Overall, individual features showed low importance in the predictions made by the model, with permutation of the most important AGE causing only a mean 1% drop in model accuracy (Figure 3B). The vast majority of features (2,979/3,013) had no impact on out-of-bag accuracy when randomly permuted (Supplementary Table 7), indicating that the machine learning model bases decisions on a genomic signature predictive of virulence level rather than on identifying individual virulence or anti-virulence factors. If a given AGE is randomly permuted, it appears that other correlated features compensate for it. Each individual AGE was included as a feature in a minority of the 10,000 decision trees, with the most prevalent AGE appearing in only 148 trees in the final model (Figure 3C). As such, it is not possible for a single AGE to have a large impact on the prediction of virulence. To further assess the apparent redundancy in our feature set, we randomly divided the 3,013 AGEs in the training set into 2, 4, and 10 subsets and evaluated the performance of random forest models built using only these subsets through nested cross-validation. We found that even when training on only a smaller subset of the accessory genomic features, model accuracy remained mostly unchanged (Supplementary Figure 3). This provides further evidence that a broad genetic fingerprint, rather than individual virulence or anti-virulence factors, is being used to classify strains as high or low virulence.

With the low permutation importance of any individual AGE, one must be cautious in drawing conclusions about their role in virulence. However, looking at the AGEs most predictive of virulence class and how they relate to one another may provide insights into genomic characteristics that are associated with, though not necessarily causative of, differences in pathogenicity. All of the ten most predictive AGEs in the random forest model were more prevalent in low virulent isolates (Table 1, Supplementary Data File 1). This is consistent with the finding that horizontally acquired genetic elements, major components of the accessory genome^6,17^, can incur a fitness cost on the host bacterium^17^. While some genomic islands encode virulence factors^11^, many horizontally acquired elements can have a parasitic relationship with the bacterium^17^. The AGE with the highest permutation importance aligns to a gene encoding for the conjugative protein TraD, perhaps suggesting a general association of conjugative elements with reduced virulence. Four of the top ten AGEs are comprised of sequences from the same “bin” in clustAGE analysis. This indicates that in at least some strains they are located near each other on the genome (i.e. part of a single, larger element). One of these four AGEs encodes an integrative and conjugative element (ICE) protein. These findings suggest that these AGEs are markers for a larger variable element common in low virulence strains. Two other AGEs are part of the same gene encoding a hypothetical protein. Finally, genes encoding for arsenic resistance are highly prevalent in low virulence isolates, perhaps suggesting either that this resistance comes at a cost or that strains adapted to survive heavy metal exposure are less able to cause disease in animals.

**Table 1.** AGEs Most Predictive of Virulence in the Accessory Genome Random Forest Model

| AGE | Mean OOB<br>Permutation<br>Importance | Subelements | Total<br>Length<br>(bp) | Total<br>Prevalence | Prevalence<br>High<br>Virulence | Prevalence<br>Low<br>Virulence | Putative<br>Annotation<sup>a</sup> |
| --- | --- | --- | --- | --- | --- | --- | --- |
| unique_grp_5582 | 0.0100 | bin364_se00006 | 433 | 0.417 | 0.161 | 0.661 | TraD |
| unique_grp_6841 | 0.0069 | bin610_se00004 | 902 | 0.304 | 0.107 | 0.492 | Hypothetical protein |
| unique_grp_1425 | 0.0063 | bin20_se00056 | 1717 | 0.330 | 0.125 | 0.525 | TetR/AcrR family<br>Transcriptional<br>regulator, Short chain<br>dehydrogenase |
| unique_grp_6842 | 0.0063 | bin610_se00005 | 369 | 0.296 | 0.089 | 0.492 | Hypothetical protein |
| unique_grp_6989 | 0.0063 | bin654_se00007 | 436 | 0.313 | 0.107 | 0.508 | Intergenic region |
| unique_grp_1437 | 0.0062 | bin20_se00073<br>bin20_se00075 | 2009 | 0.339 | 0.125 | 0.542 | SoxR, MerR family<br>DNA-binding<br>transcriptional<br>regulator, ICE<br>relaxase PFGI-1<br>class, Hypothetical<br>protein |
| unique_grp_8120 | 0.0058 | bin987_se00001<br>bin1807_se00001 | 2821 | 0.339 | 0.125 | 0.542 | AsrR family<br>transcriptional<br>regulators, Arsinic<br>transporter, Arsenate<br>reductase, ArsH,<br>Hypothetical protein |
| unique_grp_1423 | 0.0057 | bin20_se00054<br>bin20_se00057 | 1278 | 0.348 | 0.125 | 0.559 | Type II<br>glyceraldehyde-3-phosphate<br>dehydrogenase |
| unique_grp_1435 | 0.0057 | bin20_se00069 | 509 | 0.365 | 0.143 | 0.576 | Hypothetical protein |
| unique_grp_5112 | 0.0057 | bin258_se00005 | 419 | 0.357 | 0.143 | 0.559 | ArsH |

<sup>a</sup>Based on annotation of any ORF with at least 50 bp overlap with the AGE sequence when blasted against the Pseudomonas Genome Database<sup>57</sup>

### Assessing Model Performance with an Independent Test Set

The nested cross-validation performance of our random forest model provided an estimate of how well it would generalize to new *P. aeruginosa* isolates. To follow up on this, we applied the final random forest model built using all 115 training isolates to an independent test set of *P. aeruginosa* isolates to examine how well it predicted their virulence. As our test set, we selected 25 genetically diverse *P. aeruginosa* isolates previously cultured from patients with bacteremia in Spain between 2008-2009^34^ and which we have whole genome sequenced (Supplementary Table 1 and Figure 4A). The virulence of each isolate was assessed in the mouse model of bacteremia, and isolates were classified as high or low virulence using the same threshold (estimated mLD50 of 6.9 log_10_ CFU) defined for the training set (Figure 4B and Supplementary Tables 2-3). The test set was somewhat more pathogenic than the training set, with 15/25 (60%) of isolates classified as high virulence. We identified which of the 3,013 AGEs used as training features were present in each of the test isolates (Supplementary Table 8). Adding these isolates to a Bray-Curtis dissimilarity heatmap of AGE presence/absence showed that the test set is also relatively diverse in accessory genomic content (Figure 4C), a finding supported by multiple correspondence analysis (Supplementary Figure 1C-E).

**Figure 4.**
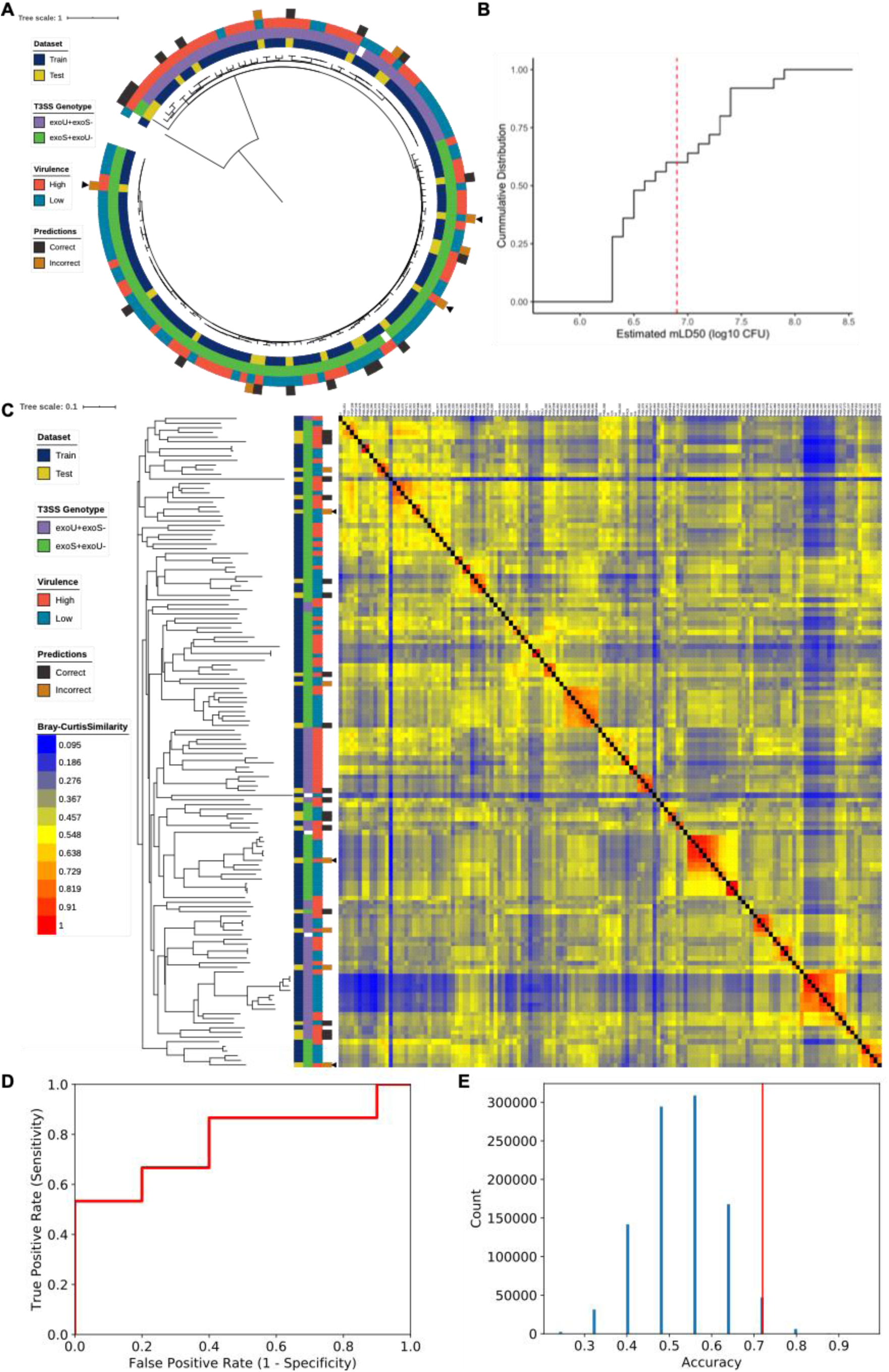
Characteristics of a random forest model trained on the accessory genomic content of the 115 *P. aeruginosa* training isolates to predict the virulence of an independent test set of 25 isolates. (A) Mid-point rooted core genome phylogenetic tree of the 115 training isolates and 25 test isolates constructed from SNV loci present in at least 95% of genomes, annotated (from inner to outer rings) with dataset, T3SS genotype, virulence level, and accuracy of prediction by the random forest model in test set. Arrowheads indicate examples of incorrectly classified test set strains whose closest core and accessory genomic neighbor(s) show a discordant virulence phenotype. (B) Cumulative density function of estimated mLD_50_ values for the 25 *P. aeruginosa* isolates making up the independent test set in a mouse model of bacteremia. Isolates with estimated mLD_50_ values less than the median estimated mLD_50_ of the training set (red dashed line) were designated as high virulence, with the remainder designated as low virulence. (C) Bray-Curtis dissimilarity heatmap comparing presence of the 3,013 AGEs identified in the training set in all 140 isolates, weighted by AGE length, and accompanying neighbor joining tree. Isolates are annotated (from left to right) by collection, T3SS genotype, virulence level, accuracy of prediction by the random forest model in test set isolates (arrowheads highlighting specific incorrectly classified test set strains as in (A)), and the dissimilarity heatmap. A higher value indicates that two isolates have more similar accessory genomes. (D) Receiver operating characteristic curve for predictions of the 25 test set isolates using the random forest model (AUC = 0.77). (E) Permutation analysis showing the likelihood of predicting test virulence with an accuracy of at least 0.72 if no true link between virulence and accessory genomic content existed. The predicted virulence of the 25 test isolates were randomly permuted 1 million times, and the resulting null distribution of possible model accuracies is shown. The vertical red line indicates the true accuracy of the random forest model in predicting test set virulence (one-sided p = 0.053).

We used the random forest model built with the training set accessory genomic and virulence information to predict the virulence of each isolate in the test set based on AGE presence or absence. Model performance on the test set (Table 2 and Figure 4D) was comparable to the estimates made through nested cross-validation. For example, the test set accuracy of 0.72 was comparable to the mean nested cross-validation accuracy of 0.75 (95% CI 0.69-0.80). This suggests that our predictive model of virulence is broadly applicable, even when tested against geographically distinct isolates. Several of the misclassified isolates in the test set appear to be exceptions in virulence when compared to their closest neighbor(s) in the core genome phylogenetic tree and the accessory genome heatmap (Figure 4A and C). Difficulty classifying these exceptional isolates is consistent with the notion that the model predictions are based on genomic signatures which perhaps approximate phylogenetic relationships. Closely related isolates that differ in virulence from the majority of their genomic neighbors would therefore be expected to be misclassified.

**Table 2.**
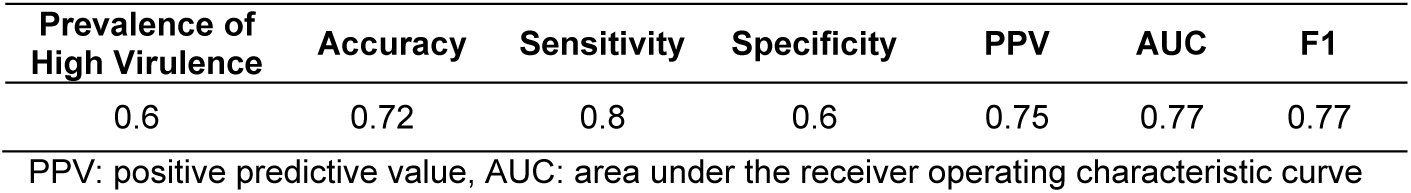
Performance of the Accessory Genome Random Forest Model Against the 25 Test Set Isolates

While it was reassuring that the random forest model performed similarly against the test set as in nested cross-validation, we wanted to ensure that the accuracy observed did not simply occur by chance. We used permutation testing to model the null distribution of test set accuracies we would expect if no link between accessory genome content and virulence existed in the test set. Using this approach, we determined the probability of observing a test set accuracy of at least 0.72. Randomly permutating the predicted virulence of the 25 test isolates one million times resulted in an accuracy of at least 0.72 in 53,476 cases (one-sided p = 0.053) (Figure 4E). The test set performance observed is therefore unlikely if the accessory genome does not predict virulence. Limiting factors include the small sample size of the independent test set, as is evident from the discrete possible accuracies when the predictions were permuted, and that we would not expect the model to perform better against new data than it did during nested cross-validation.

### Addressing Model Limitations by Removing Isolates with Intermediate Levels of Virulence

While the models generated thus far show that the accessory genome is predictive of *P. aeruginosa* virulence in mice, limitations inherent to our binary classification of virulence may constrain their performance. The first lies in the resolution of the mLD_50_ estimates used as the basis for these classes. Because of the practical limitations of testing over 100 isolates in mice, many isolates were tested with only two or three doses. This leads to uncertainty in the dose required to cause severe disease (Supplementary Tables 2-3). Second, isolates with mLD_50_ estimates close to the cutoff may actually be quite similar, both in their virulence and in their genomic makeup, but still be assigned to different virulence classes. To assess the extent to which this ambiguity influenced the results, we repeated the machine learning pipeline using the random forest algorithm after removing intermediate virulence isolates (the middle third of estimated mLD_50_ values). This enforced a greater separation of isolates classified as high and low virulence (Figure 5A). Even with a third fewer training isolates, nested cross-validation performance was similar to when all training isolates were included, with a mean accuracy of 0.76 (95% CI 0.67-0.85) (Figure 5B). The learning curve, however, showed a greater distance between the training and cross-validation scores (Figure 5C). This suggests a higher potential performance when intermediate virulence isolates are removed. The benefit of having a clearer boundary between high and low virulence would likely become apparent with a larger training set, though the number needed and degree of improvement is unclear.

**Figure 5.**
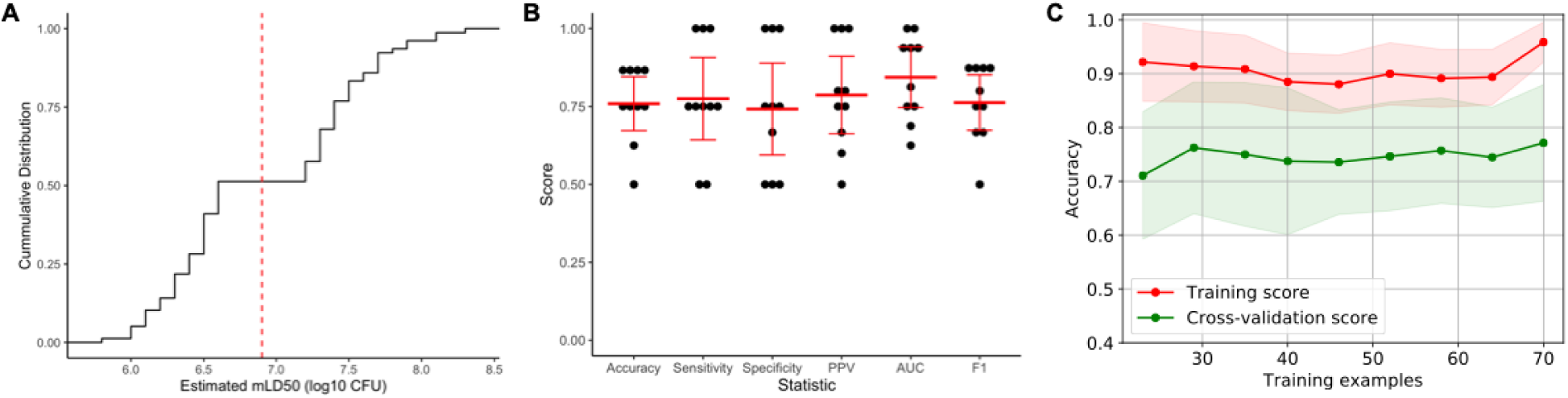
Performance of the random forest algorithm in predicting *P. aeruginosa* virulence from accessory genomic content when intermediate virulence isolates (middle 3^rd^ of estimated mLD_50_s) were removed. (A) Cumulative density function of estimated mLD_50_ values after removing intermediate virulence isolates. Isolates with estimated mLD_50_ values less than the median value in the complete training set (red dashed line) were designated as high virulence, with the remainder designated as low virulence. (B) Nested 10-fold cross-validation performance of the random forest model, including accuracy, sensitivity, specificity, positive predictive value (PPV), area under the receiver operating characteristic curve (AUC), and F1 score. The results for each cross-validation fold are shown in black with the mean and 95% confidence interval of each statistic indicated in red. (C) Learning curve showing change in mean training accuracy (red line) and cross validation accuracy (green line) with increasing training set sizes. Shading indicates the 95% confidence interval. Assessments at each number of training examples were through 10-fold nested cross-validation.

### Incorporating Test Set Isolates into the Accessory Genome Model

After using the 25 additional isolates as an independent test set, we next examined their impact on nested cross-validation performance if they were included in the training set. As this changes the median estimated mLD_50_, we performed the modeling using both the median of the 115 training set isolates and the median of all 140 isolates as the cutoff for high/low virulence. These models performed similarly, both to each other and to the results seen with only the original training set. The mean nested cross-validation accuracy was 0.72 (95% CI 0.65-0.79) when using the median mLD_50_ cutoff of the 115 training isolates and 0.69 (95% CI 0.60-0.78) when using the median mLD_50_ cutoff of all 140 isolates (Supplementary Figure 4C and E). The learning curves, however, showed greater overfitting of the model when the all-isolates median cutoff was used, with a larger separation between the training and cross-validation accuracies (Supplementary Figure 4D and F). This suggests the choice of cutoff between high and low virulence isolates may become more important with increasing training set sizes. Removing intermediate virulence isolates resulted in similar nested cross-validation performance and learning curves as seen when performing this analysis on the original training isolates (Supplementary Figure 4B, G, and H).

### Modeling *P. aeruginosa* Virulence with Features Incorporating Core Genome Information

Thus far we have shown that the accessory genome of *P. aeruginosa* is predictive of strain virulence. The accessory genome and core genome are correlated with each other, as can be seen from previous reports^4^ and by comparing core and accessory genome measures of strain relatedness (Figure 1B and C). As such, the accessory genome contains implicit information about the core genome. Still, it is possible that our focus on the accessory genome misses important core features predictive of virulence. To address this possibility, we defined our feature set in two additional ways and examined the performance of random forest models trained using these features. First, we considered core genome SNVs, using one-hot encoding in our machine learning pipeline to convert the SNVs into binary variables interpretable by the algorithm. Second, we used whole genome k-mer counts, which encode information about variability in both the accessory and core genome, considering a k-mer length of both 8 and 10 bp. Unlike the AGE feature set used previously, which considered the presence and absence of accessory elements, the k-mer feature sets additionally capture polymorphisms within these elements. We estimated the performance of approaches using these feature sets through nested cross-validation and then assessed how well final models built with each were able to predict the virulence of the 25 independent test set isolates.

A random forest approach using core genome SNVs as features performed worse on average in nested cross-validation than when using accessory genomic features, with a mean accuracy of 0.65. However, its 95% confidence interval (0.55-0.75) still overlapped with those seen for the accessory genomic models (Figure 6A). Therefore, some information important for determining virulence level may be missed by not considering the accessory genome. Another explanation is that more strains may be needed to model this substantially more complex feature set, as there were 440,116 core genome SNV loci detected in our training set. As the confidence intervals overlap, we must be careful drawing conclusions about the relative predictive power of the core and accessory genomes. The final model trained with core genome SNV features showed an accuracy of 0.72 on the independent test set (Supplementary Table 9). This was identical to the test set accuracy seen for the accessory genomic model but was associated with a lower sensitivity and higher specificity (Table 2 and Supplementary Table 9). Despite its lower nested cross-validation accuracy, we therefore cannot say whether the accessory genome or core genome are superior in predicting virulence.

**Figure 6.**
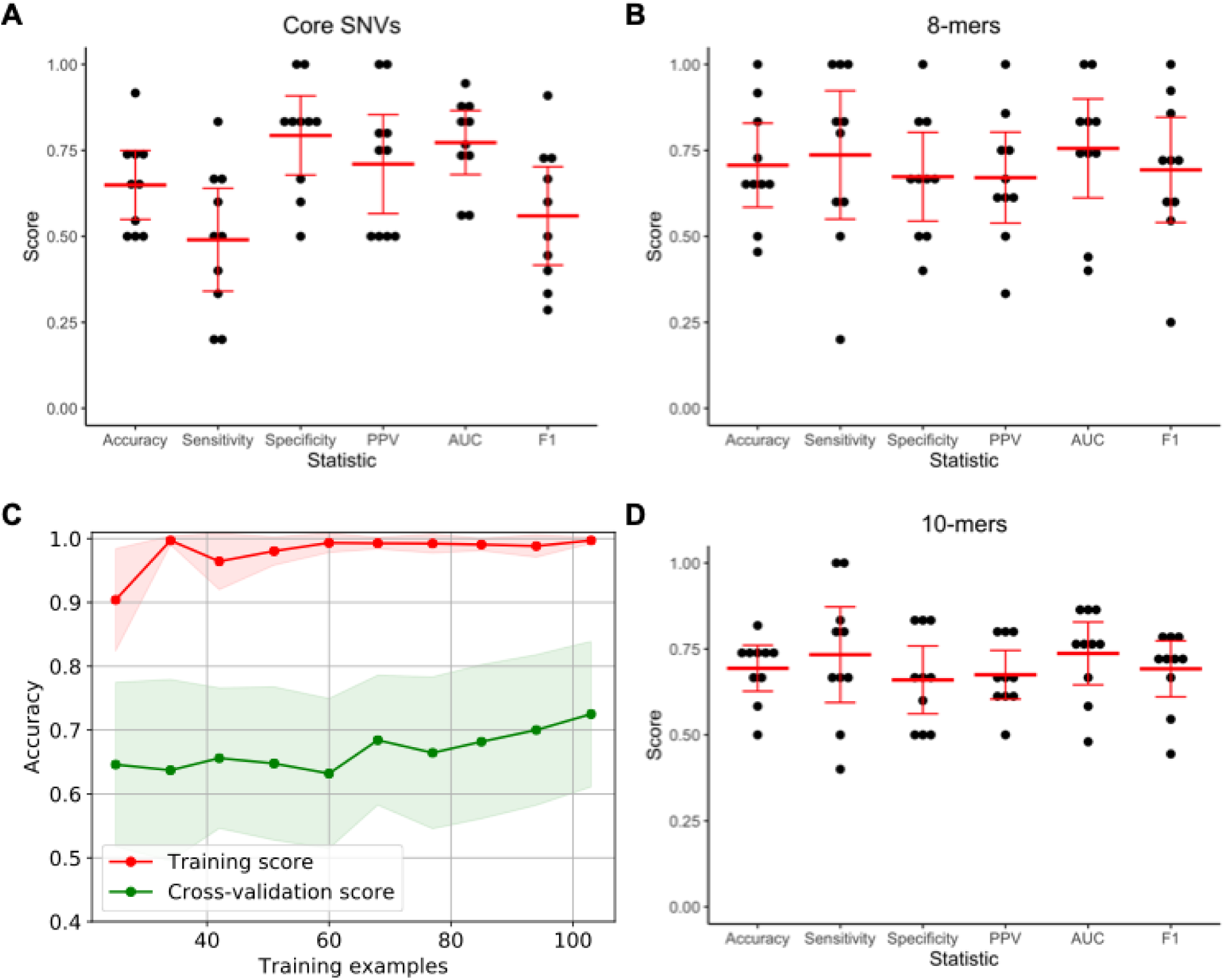
Performance of the random forest algorithm in predicting *P. aeruginosa* virulence when 8-mer counts, 10-mer counts, or core genome SNVs were used as model features. Nested cross-validation performance when using (A) core genome SNVs, (B) 8-mer counts, and (D) 10-mer counts, including accuracy, sensitivity, specificity, positive predictive value (PPV), area under the receiver operating characteristic curve (AUC), and F1 score. The results for each cross-validation fold are shown in black with the mean and 95% confidence interval of each statistic indicated in red. (C) Learning curve showing change in mean training accuracy (red line) and cross validation accuracy (green line) when using 8-mer counts as features as increasing numbers of isolates are used to train the random forest model. Shading indicates the 95% confidence interval. Assessments at each number of training examples were through 10-fold nested cross-validation. Learning curves were not constructed when using core genome SNV or 10-mer counts as features for reasons of computational feasibility.

The random forest approach using k-mer counts as features performed similarly to the accessory genome models in nested cross-validation, with a nested cross-validation accuracy of 0.71 (95% CI 0.58-0.83) when 8-mer counts were used and 0.69 (95% CI 0.63-0.76) when 10-mer counts were used (Figure 6B and D). This suggests that no additional predictive information was gained from incorporating core genome features, and that AGE presence/absence encodes the same information in a smaller feature set. The learning curve for the model trained on 8-mer counts showed overfitting, with a large discrepancy between the training and cross-validation accuracies (Figure 6C). This suggests that performance would improve with a larger training set, and perhaps that the increased complexity of the 8-mer feature set makes it more difficult to learn from than the presence or absence of AGEs. The final model trained with 8-mer features showed an accuracy of 0.60 on the test set, while the final model trained on the 10-mer feature set showed an accuracy of 0.68 (Supplementary Table 9). The performance of the 8-mer feature set was more variable in nested cross-validation, with a wider range in its 95% confidence interval, and it is possible that lower model stability contributed to its poorer performance against the test set.

## DISCUSSION

In this study, we have shown that a signal exists in the *P. aeruginosa* accessory genome that is predictive of an isolate’s virulence in a mouse model of infection. This finding was consistent across a variety of machine learning algorithms. Results for the random forest approach were validated using an independent test set of clinical isolates collected from a geographically distinct source, showing the broad applicability of the *P. aeruginosa* accessory genome in predicting virulence. We additionally showed that the core genome, alone or in combination with the accessory genome, is also predictive of virulence, but the ability of models trained on this information to generalize to the independent test set was less conclusive. These types of genetic features were substantially more complex, and models trained from them may benefit from increasing sample size. The machine learning analyses conducted here serve as a framework to further investigate the relationship between the genome of a bacterium and its phenotype.

The random forest model trained on accessory genomic information classified isolates as high or low virulence based on a diffuse genomic signature rather than by detecting a small number of virulence or anti-virulence factors. These results suggest that lineage may predict phenotype, echoing the recent finding that genomic neighbors are highly predictive of antimicrobial resistance in *Streptococcus pneumoniae* and *Neisseria gonorrhoeae*^*26*^. As such the genomic signature detected may approximate lineage. These findings also highlight the complexity of *P. aeruginosa* virulence compared to antimicrobial resistance, which is often dictated by one or a few genes or mutations. Still, it is relevant that all of the ten most important AGEs included in our model were associated with low virulence. This supports the earlier finding that *P. aeruginosa* accessory genes can reduce virulence in *C. elegans* and that active CRISPR systems, which would limit acquisition of foreign DNA and new AGEs, are associated with higher virulence in that model^16^. While certain AGEs enhance virulence^15^, the majority of AGEs (e.g. parasitic phages, plasmids, or ICEs) may decrease virulence. We focused on virulence in a mouse model of acute infection, and as such certain bacterial genetic factors important in the hospital setting may not apply. Antimicrobial resistance, for example, can be an important prognostic factor for patient outcomes^35,36^ but would not be relevant in this model. Future studies should examine the types of AGEs that are associated with, and ultimately causal of, both increased and decreased virulence and how this varies between infection models.

Our random forest model built on accessory genomic features showed similar performance in nested cross-validation as when applied to an independent test set of 25 isolates. By looking at the test set isolates that were classified incorrectly, we can learn why the model sometimes failed. Some incorrect predictions may be because of mLD50s near the threshold between high and low virulence, leading to ambiguity in their true virulence level. An example of this scenario is the isolate PASP518, whose estimated mLD_50_ of 7.0 log_10_ CFU is near the cutoff of less than 6.9 log_10_ CFU for high virulence. This highlights an inherent limitation of this study, in that virulence exists on a continuum not neatly divided into binary classes. To address this limitation, we examined how the model performs when excluding intermediate virulence isolates. In this condition, a random forest approach performed similarly in nested cross-validation with a third less samples and learning curve analysis showed a potential for higher accuracy with increasing sample size (Figure 5). On the other hand, some of the incorrect predictions in the test set were exceptions in virulence compared to closely related isolates. For example, PASP251 has an estimated mLD_50_ of 6.3 log_10_ CFU, while its nearest four phylogenetic neighbors all have estimated mLD_50_s of greater than 7 log_10_ CFU. This could be because PASP251 possesses additional virulence determinants in the form extra accessory genes or distinct core genome polymorphisms. In either case, PASP251 is a particularly interesting isolate for future study. An alternative explanation is that, while geography does not seem to play a role in model performance as a whole, the closely related isolates from Chicago have acquired common mutations or genes modifying their virulence. An increased sample size may ameliorate the problem of isolates being misclassified by allowing for finer resolution of subgroups that are associated with high or low virulence, especially if the model were able to learn new and more discriminatory patterns of features. Learning curve analysis for the random forest approach (Figure 3A) suggests the impact of adding more isolates would be limited, but this cannot account for new or more predictive features that could arise from increasing the amount of genetic data available.

As whole genome sequencing becomes an increasingly routine component of clinical microbiology practice, it will create the opportunity to risk-stratify patients based on the genome of an infecting bacteria and influence treatment decisions in real-time. The ability of the genome to predict antibiotic resistance has been established^20,21,23,25,26^, opening the door for sequencing to supplement or replace traditional antimicrobial susceptibility testing. This study serves as a proof of concept that the *P. aeruginosa* genome can be used to predict its pathogenicity. Future studies are needed to expand beyond virulence in mice and provide a more complete understanding of the role genetic variation plays the ability of *P. aeruginosa* to cause disease. An area of particular interest is in predicting patient outcomes from the genome of an infecting isolate. Large retrospective studies using archived isolates with corresponding clinical data would allow for exploration of the relative importance that bacterial and patient factors play in predicting patient outcomes, as has been shown for specific *S. aureus* clones^28^. This could improve the sophistication of current diagnostics and allow clinicians to rapidly identify patients at highest risk for poor outcomes.

## METHODS

### Bacterial Isolates

A training set of *P. aeruginosa* isolates for use in the machine learning analyses was established as follows. A total of 98 isolates previously collected at NMH in Chicago, USA from 1999-2003 from adults with *P. aeruginosa* bacteremia^31^ were selected after exclusion of 2 isolates that had been collected from patients with a history of cystic fibrosis. An additional 17 isolates from pediatric patients with Shanghai Fever collected at Chang Gung Children’s Hospital in Taiwan from 2003-2008^32^ were included. This yielded a training set size of 115 isolates. A genetically diverse independent test set of 25 isolates was selected from a larger cohort of isolates collected from patients with bacteremia in Spain between 2008-2009^34^ (Supplementary Table 1).

### Mouse Model of Bacteremia

Female 6- to 9-week-old BALB/c mice were infected via tail-vein-injection in a model of bacteremia as previously described^32^. Isolates were plated from freezer stocks onto lysogeny broth (LB) agar, and single colonies were inoculated into MINS broth^37^ and grown overnight at 37 °C. Overnight cultures were then subcultured in fresh MINS broth for approximately 3 hours at 37 °C. Cultures were resuspended in PBS before dilution to the target dose, and 50 μL was injected into each mouse via the tail vein. Inocula, in CFUs, were then determined by serial dilution, plating, and colony counts. Mice were monitored for the development of severe disease over 5 days, with mice exhibiting endpoint disease euthanized and scored as dead. Each isolate was tested at a minimum of 2 doses, with 3-5 mice per dose (minimum 9 total mice per isolate) (Supplementary Table 2). Many of the mouse experiments included in this study were previously reported as part of other studies. In particular, the majority of experiments with the NMH strains were performed as part of Allen *et al.*, 2020^15^. Some experiments with the Taiwan isolates PAC1 and PAC6 were performed as part of Chuang *et al.*, 2014^32^.

A modified 50% lethal dose (mLD_50_) for each isolate was estimated from the above experiments using the drc package (v3.0-1)^38^ in R (v3.6.1)^39^. One outlier experiment for strain S2, which caused 20% mortality at a dose of ∼7.2 log_10_ CFU, was excluded as doses of ∼6.3 and ∼6.8 log_10_ CFU caused 80% and 100% mortality, respectively, in other experiments. Percent mortality as a function of dose (in units of log_10_ CFU) was modeled using a two-parameter log-logistic function and binomial data type. These models were used to estimate the mLD_50_ for each isolate, which was then rounded to the nearest tenth (Supplementary Table 3). Isolates with rounded mLD_50_ estimates below the median were classified as high virulence, with the remainder classified as low virulence.

All experiments were approved by the Northwestern University Institutional Animal Care and Use Committee in compliance with all relevant ethical regulations for animal testing and research.

### Whole genome Sequencing and Assembly

Short-read whole genome sequencing was performed for all isolates using either Illumina HiSeq or Miseq platforms to generate paired-end reads. Reads were trimmed using Trimmomatic (v0.36)^40^, with Nextera adapter removal, a sliding window size of 4 bp with average quality threshold of 15, and a minimum trimmed read length of 36 bp. Draft genomes were assembled from trimmed paired-end reads using SPAdes (v3.9.1)^41^ with the careful and automatic read coverage cutoff options. Draft genomes were further filtered to remove contigs shorter than 200 bp, with less than 5-fold mean read coverage, or with alignment to phiX. Even using only trimmed reads, the mean coverage of each filtered assembly was at least 24-fold.

Many of the whole genome sequences used in this study were previously reported as parts of other studies^15,42,43^. Draft genomes originally assembled through different methodologies were re-assembled as described above.

For several genomes (PABL012, PABL017, PABL048, PAC1, and PAC6), long-read sequencing and hybrid assembly was performed. Briefly, genomes were sequenced on the PacBio RS II platform. Raw data were assembled using the HGAP assembler (SMRT Analysis v2.3.0), Canu assembler (v1.2)^44^, and Celera assembler (v8.2)^45^, all using default settings. Contigs were combined and circularized using Circlator (v1.5.1)^46^. Assemblies were polished using Quiver (SMRT Analysis v2.3.0). Indel errors were corrected using Pilon (v1.21)^47^ using paired-end reads generated on Illumina HiSeq or MiSeq platforms. The complete genome for PABL048 was generated as part of a previous study^42^.

### Phylogenetic Analysis

kSNP (v3.0.21) was used to generate 95% core genome parsimony phylogenetic trees for both 115 isolates in the training set and all 140 isolates in the training and test sets, using fasta files as input. The Kchooser program was used to select the optimum k-mer size of 21, and SNP loci present in at least 95% of input genomes were used to make the trees^48^. The phylogenetic trees were annotated and plots generated using iTOL (v4)^49^.

### Accessory Genome Determination

Accessory genomes for the 115 *P. aeruginosa* isolates in the training set were determined using the programs Spine (v0.3.2), AGEnt (v0.3.1), and ClustAGE (v0.8)^7,50^. Spine was used with Prokka^51^-annotated genbank files for each isolate as input to generate a core genome of sequences present in at least 95% of isolates. AGEnt was then used to determine the accessory genome of each isolate based on comparison to the core genome. The accessory genomes of all 115 isolates were then compared using ClustAGE to identify shared sequences using an 85% identity cutoff. ClustAGE identifies the longest continuous accessory sequences as “bins” and the portions of these bins that differ from isolate to isolate as “subelements”^15,50^. As part of this process, the read correction feature of ClustAGE was used to identify sequences present in the original sequencing reads that were missed during genome assembly. All perfectly correlated subelements identified through clustAGE were collapsed into a single feature, termed a “unique group (of subelements)”. For the purpose of this study, accessory genomic elements (AGEs) were defined as all unique groups totaling ≥ 200 bp. A dataframe of all AGEs in the training isolates served as the accessory genome feature set in subsequent machine learning analyses (Supplementary Tables 4-5). To generate AGE features present in all genomes (both the original training and test sets), this process was repeated using all 140 *P. aeruginosa* isolates as input (Supplementary Table 10).

To determine which AGEs from the training set were present in the test set, clustAGE was run using the training set read-corrected subelement sequences (for all subelements ≥ 50 bp) from the 115 training isolates as a reference AGE set with the “--AGE” option and comparing to the draft genomes of all isolates in the test set, with read correction to identify any sequences present that were not included in draft genome assembly. This identified which portions of each subelement were found in the test set with an 85% identity cutoff. An AGE (defined as a unique group of subelements) was called as present if at least 85% of the screened length was detected (Supplementary Table 8).

To examine the relationships between accessory genomes in the training isolates, their AGE content was compared using the subelement_to_tree.pl utility from ClustAGE. This calculated the Bray-Curtis dissimilarity between each isolate based on AGE presence or absence, with the impact of each AGE weighted by its length. A neighbor joining tree was constructed from 1000 bootstrap replicates using the matrix of Bray-Curtis dissimilarities. For consistency with the definition of AGE used in this study, unique groups of subelements were used as input. The neighbor joining tree and associated heatmap of Bray-Curtis dissimilarities were annotated and visualized with iTOL (v4)^49^. To examine the accessory genomic relatedness of the 25 test set isolates based on training-set derived AGEs, the training set AGE calls defined above were added and Bray-Curtis dissimilarity calculations, and neighbor joining tree construction were repeated. To further evaluate the relationships between accessory genomes, multiple correspondence analysis (MCA) was performed based on the presence or absence of AGEs in the 115 training isolates. Additionally, MCA was perfumed considering which of the training isolate AGEs were identified all 140 isolates. MCA was performed in R (v3.6.1)^39^ using the FactoMineR (v2.3)^52^ package and visualized using the factoextra (v1.0.6) package.

### Sequence Alignment and Core SNV Calling

Sequence alignment of paired-end Illumina reads for each genome to the reference genome PAO1 (RefSeq accession NC_002516) was performed as previously described^42^. Briefly, reads were trimmed with Trimmomatic (v0.36)^40^ and aligned to PAO1 with BWA (v0.7.15)^53^. Loci passing inclusion criteria were called as having the PAO1 base or a SNV base for each genomic position, with the remainder of positions converted to gaps. PAO1 alignments for all 115 training isolates were concatenated, SNV positions present in fewer than 95% of genomes were filtered, and invariant sites were then removed. This core variant SNV alignment was used as the SNV feature set in subsequent machine learning analyses, with a one-hot-encoding step added to the pipeline to convert SNV loci into multiple binary variables. This feature set was defined in the test set by considering the genomic positions identified as variant in the training set. By extracting the sequence present at these variant positions in the PAO1 alignments for each of the 25 test set isolates, we created a SNV feature set corresponding to that used in the training set.

### K-mer Counts

K-mer counts (using either 8 or 10 bp k-mers) were determined for each genome using KMC3 (v3.0.0)^54^. All k-mers occurring at least once in each genome’s fasta file were identified using the kmc application, and a count file was generated using the kmc_dump application. All unique k-mers identified in the training set of 115 *P.* aeruginosa genomes were used to construct a dataframe of k-mer counts for each genome. This served as k-mer feature set in subsequent machine learning analyses. This feature set was defined in the 25 test set isolates by considering the counts of all k-mers previously identified in the training set.

### Predicting Virulence Based on Genomic Features

Machine learning analyses were performed using the sci-kit learn library (v0.21.2)^55^ in Python (v3.6.9). The general workflow for the machine learning pipeline is described in Supplementary Figure 2. A training dataset of features (AGEs, k-mers, or core SNVs) and labels (high/low virulence) was defined. A machine learning algorithm (random forest, l2-regularized logistic regression, elastic net logistic regression, or support vector classifier) was chosen, and a grid of relevant hyperparameters to test were defined. A machine learning model was then trained using the selected algorithm, with hyperparameter tuning performed through grid-search cross-validation. A 10-fold stratified cross-validation strategy was used. This generated a final model which can be used to predict the virulence class of new isolates. Concurrently the generalization performance of this model was estimated through nested cross-validation. In this process, grid-search cross-validation was performed within an outer 10-fold stratified cross-validation loop. The performance of a grid-search cross-validation tuned model against each cross-validation fold was determined (including accuracy, sensitivity, specificity, positive predictive value, area under the receiver operating characteristic curve, and F1 score). The median, mean, and 95% confidence interval of the nested cross-validation results were determined and plotted with the values for each fold using R (v3.6.1)^39^ with the tidyverse library suite (v1.2.1)^56^.

For the random forest algorithm, the number of trees was set to 10,000 and “max_features”, “min_samples_split”, “min_samples_leaf”, “criterion”, and “max_depth” were varied as hyperparameters during grid-search cross-validation. The logistic regression algorithm was considered using l2 regularization (penalty = “l2”) and elastic net regularization (penalty = “elasticnet”) separately. For l2-regularized logistic regression, the “lbfgs” solver was used, “max_iter” was set to 10,000, and “C” was varied as a hyperparameter during grid-search cross-validation. For elastic net logistic regression, the “saga” solver was used, “max_iter” was set to 10,000, and “C” and “l1_ratio” were varied as hyperparameters. For the support vector classifier algorithm, the radial basis function kernel was used, and “C” and “gamma” were varied as hyperparameters during grid-search cross-validation.

In some cases, learning curves were created to examine how training and nested cross-validation accuracy varied with increasing training test size. For this, the dataset was split into training and cross-validation folds through 10-fold stratified cross-validation. Subsets of examples were then drawn from each training fold ranging from 25% to 100% of the training fold size. On each subset, a model was trained through the grid-search cross-validation approach described above. The mean and 95% confidence interval for training and cross-validation accuracies at each number of examples were then determined and plotted.

### Random Forest Permutation Importance

Out-of-bag permutation importance for the random forest model of virulence based on accessory genomic content trained on the complete training set of 115 *P. aeruginosa* isolates was determined using the rfpimp (v1.3.4) Python package (https://github.com/parrt/random-forest-importances). This measures the decrease in accuracy in predicting out-of-bag samples (samples not used to train a given decision tree in the random forest) if a feature is randomly permuted. As the impact of permuting a given feature on model accuracy may depend on how it is permuted, this process was repeated a total of 100 times to determine a mean permutation importance (Supplementary Table 7). The putative annotation of the top 10 AGEs identified by permutation importance was determined by blast search of subelement sequences against the Pseudomonas Genome Database^57^ and including the annotation of any ORF for which at least 50 bp were contained in the AGE.

### Evaluating Random Forest Model Performance with an Independent Test Set

The random forest model trained on AGE presence/absence in the 115 training isolates were tested against the independent test set of 25 isolates. The training-set AGEs identified in these 25 isolates were used as features, and the predicted virulence classes were compared to the actual virulence for these isolates. This was used to estimate testing accuracy, sensitivity, specificity, positive predictive value, area under the receiver operating characteristic curve, and F1 score and to plot the receiver operating characteristic curve. This approach was also used to assess the performance of random forest models trained on core genome SNVs, 8-mers, and 10-mers against the independent test set of 25 isolates.

For the accessory genome model, the probability of seeing the observed test set accuracy by chance if there were no true association between the predicted virulence (and therefore accessory genome) of an isolate and its true virulence was estimated through permutation testing. The predicted virulence classes for the 25 test isolates were randomly permuted 1 million times and used to create a null distribution of possible model accuracies. The observed test set accuracy was compared to this null distribution to estimate a one-sided p value.

## DATA AVAILIBILITY

BioSample accessions for all isolates used in this study are listed in Supplementary Table 1. For all isolates, the version of the genome assemblies used in this study are available on GitHub. Input data for machine learning analyses are also available on GitHub (https://github.com/nathanpincus/PA_Virulence_Prediction).

## CODE AVAILIBILITY

Code used for machine learning analyses in this study, including details on hyperparameters used during grid-search cross-validation, and for plotting the results are available on GitHub (https://github.com/nathanpincus/PA_Virulence_Prediction).

## ACKNOWLEGEMENTS

This work was supported by the National Institute of General Medical Sciences [T32 GM008061 and T32 GM008152 (N.B.P)], the American Cancer Society [MRSG-13-220-01-MPC (E.A.O)], and the National Institute of Allergy and Infectious Diseases [R01 AI118257, R21 129167, K24 AI104831, and U19 AI135964 (A.R.H.)]. J.J.D. and M.N. are supported by the United States Defense Advanced Research Projects Agency Friend or Foe program iSENTRY award [Contract No. HR0011937807 (J.J.D.)], and by the United States National Institute of Allergy and Infectious Diseases Bacterial and Viral Bioinformatics Resource Center award [Contract No. 75N93019C00076 (PI Rick Stevens)]. A.O. is supported by Instituto de Salud Carlos III, Subdirección General de Redes y Centros de Investigación Cooperativa, Ministerio de Economía y Competitividad, Spanish Network for Research in Infectious Diseases (REIPI RD16/0016/0004), co-financed by European Development Regional Fund “A way to achieve Europe” and operative program Intelligent Growth 2014-2020. This research was supported in part through the computational resources and staff contributions provided by the Genomics Compute Cluster, which is jointly supported by the Feinberg School of Medicine, the Center for Genetic Medicine, and Feinberg’s Department of Biochemistry and Molecular Genetics, the Office of the Provost, the Office for Research, and Northwestern Information Technology. The Genomics Compute Cluster is part of Quest, Northwestern University’s high-performance computing facility, with the purpose to advance research in genomics. We acknowledge the University of Maryland School of Medicine Institute for Genome Sciences for performance of PacBio whole-genome sequencing.

## AUTHOR CONTRIBUTIONS

N.B.P. and A.R.H. conceived the study. N.B.P., A.R.H, J.J.D., M.N., and D.R.W. designed the experiments. N.B.P and E.A.O. performed bioinformatic analyses. N.B.P. performed computational experiments. N.B.P., E.A.O., and J.P.A. performed *in vivo* experiments. C.-H. Chuang, C.-H. Chiu, L.Z., and A.O. provided instrumental resources. All authors provided intellectual contributions and reviewed the paper.

## MATERIALS AND CORRESPONDENCE

Correspondence and requests for materials should be addressed to A.R.H.

